# Mecp2 Deficiency Alters M1/M2 Gene Expresion in Bone Marrow-Derived Macrophages Upon Stimulation

**DOI:** 10.1101/2020.08.18.256313

**Authors:** M.I Zalosnik, B De Filippis, R. De Simone, D. Pietraforte, G. Laviola, A.L Degano

## Abstract

Rett Syndrome (RTT) is a neurodevelopmental disorder mostly caused by mutations in the X-linked gene, MeCP2, which encodes for methyl-CpG binding protein 2 (MeCP2). MeCP2 is member of a family of methyl binding proteins that control the expression of several genes according to the genomic context. Emerging evidence suggests that immune dysfunctions would actively contribute to the pathogenesis of RTT. Macrophages are key effector cells that participate in several critical aspects of immune responses. The aim of our work was to assess the response of macrophages *in vitro* in the context of polarizing stimuli. We used bone marrow-derived macrophages (BMDM) obtained from MeCP2^308/y^ mice, a mouse model that carries a truncated form of MeCP2. Since MeCP2 is expressed as a “partially functional” protein in humans with RTT it becomes crucial to establish how the presence of a mutant form of MeCP2 affects immune responses to support the normal homeostasis of individuals. MeCP2 deficiency induced exacerbation of pro-inflammatory mediators and deficient immune regulatory responses under polarizing conditions. These findings suggest that MeCP2 plays a role in the establishment of macrophage polarization in the context of immune activation. Present results may have important implications in understanding RTT pathogenesis and for developing potential treatments.

**Conflict of interest:** The authors declare no conflicts of interest.

## INTRODUCTION

Rett Syndrome (RTT, OMIM #312750) is an X-linked neurological disorder characterized by an apparent normal development until 6 to 18 months of age followed by a progressive loss of motor skills, accompanied with cognitive and social disabilities. The incidence is ~1/10,000–1/15,000 female births and it is not hereditary. *De novo* mutations in the X-linked gene, MeCP2, which codes for methyl-CpG binding protein 2 (MeCP2), account for about 95% of RTT cases^1^. MeCP2 is part of a basic nuclear protein family that binds to methylated cytosine at CpG islands, which are the canonical DNA methylation sites controlling the expression of several genes according to the genomic context.

It has been proposed that dysfunctions in myeloid cells would actively contribute to the pathogenesis of RTT. So far, these studies focused mainly on microglial cells due to their close interaction with neurons. Although microglial cells and tissue-resident macrophages share a common embryonic origin^2^, MeCP2 was shown to act in a cell-dependent manner in myeloid cell populations^3^. In addition, most studies conducted to determine the function of MeCP2 in myeloid cells have used KO or *null* mouse models characterized by a total absence of protein. Since MeCP2 is expressed as a “partially functional” protein in humans with RTT^4^, we reasoned it is crucial to establish how the presence of a mutant form of MeCP2 affects immune responses to support the normal homeostasis of individuals.

Therefore, the aim of our work was to evaluate the response of macrophages in the context of different polarizing stimuli by using bone marrow-derived macrophages (BMDM) isolated from MeCP2^308/y^ mice, a mouse model that expresses a truncated form of MeCP2. Our results showed that MeCP2 mutation induces exacerbation of macrophage pro-inflammatory responses. Moreover, this is the first report demonstrating that MeCP2 modulates IL-4-induced immune response in macrophages. These results have important implications for exploring the consequences of MeCP2 mutations on immune homeostasis.

## MATERIALS AND METHODS

### Animals

We used the Mecp2^308/y^ mice which carry a premature stop codon at amino acid 308 and as result, generating a truncated MeCP2 protein that lacks the C-terminus (B6.129S-Mecp2 /J, Stock 005439, The Jackson Labs)^28^. All the experiments were performed using only hemizygous MeCP2 males (MeCP2 MUT) and their corresponding WT male littermates as control, in order to avoid the high variability caused by random X-inactivation in females. At weaning, mice were housed in groups of 2-3 in polycarbonate transparent cages (33×13×14 cm) with sawdust bedding and kept on a 12h light-dark schedule (lights off at 8:00 am). Temperature was maintained at 21 ± 1°C and relative humidity at 60 ± 10%. Animals were provided *ad libitum* with tap water and a complete pellet diet (Altromin, Germany). All procedures were carried out in accordance with the European Communities Council Directive (10/63/EU) as well as Italian law (26/2014).

### Establishment of bone marrow-derived macrophages (BMDM)

6-12 weeks of age WT and MeCP2 MUT mice were sacrificed by cervical dislocation and rapidly proceeded to obtain the bones of both hind legs in order to isolate the bone marrow of the femur and tibia. In 10 cm petri dishes, 2×10^5^ cells were cultured in 10 mL of RPMI 1640 medium supplemented with 10% endotoxin-free fetal bovine serum, 50 g/mL of Gentamicin, 2mM of L-Glutamine and 20ng/mL of macrophage colony stimulating factor (M-CSF). The cells were incubated at 37°C with 5% CO_2_. After 3 days of incubation, 10 ml of medium containing M-CSF was added. On day 6 of *in vitro* differentiation, cells were harvested. The number and viability of cells were estimated by counting 0.2% Trypan Blue solution in a Neubauer chamber.

### BMDM culture stimulations

After the harvest of the BMDM on day 6 of differentiation, cells were placed in multiwell plates for 24 h with complete RPMI to let the macrophages adhere to the plate. After, new RPMI was added and BMDM were classically activated (M1 macrophages) using LPS (100 ng/mL) and IFN_γ_ (50ng/mL). BMDM were alternatively activated by incubating them to IL-4 (10 ng/mL)). The stimulation was performed for 4 h for the experiments aimed to obtained gene expression profile analyzed by Real Time RT-PCR and for 22 h for the nitric oxide and superoxide production assays.

### Flow Cytometry

After 6 days of differentiation, BMDM were labeled with APC anti-mouse/human CD11b (1:400, Cat. 101211, BioLegend CA, USA) and PE anti-mouse CD45 (1:400, Cat. 103105, BioLegend CA, USA) and the percentage of CD11b+/CD45+ cells was determined by flow cytometry. BMDM were harvested and 1 million cells were resuspended into a final volume of 500 μl with FACS. They were incubated on ice for 30 min with anti-CD11b and anti-CD45 antibodies. After, the cells were washed with PBS and resuspended in 100 μl of PBS. At least 50,000 events were acquired from each sample using a FACS Canto II Cytometer (Becton Dickinson, San Jose, CA, USA) and analyzed with FlowJo software version 5.7.2.

### Real Time RT-PCR

Gene expression analysis was performed by real time RT-PCR. *RNA extraction.* 1×10^6^ BMDM were homogenized and resuspended in 1 ml of TRIzol reagent and proceeded according to manufacturer’s instructions. *Reverse Transcription*. 2 μg of RNA was incubated at room temperature for 15 min with DNase I. The product was incubated with random hexamer primers, deoxynucleotides and the reverse transcriptase M-MLV. Reverse transcription was performed following the manufacturer’s specifications, using a thermocycler Mastercycler gradient (Eppendorf, Hamburg, Germany) in one cycle as follows: 6 min at 25 °C, 60 min at 37 °C, 18 min at 70 °C and 10 min at 4 °C. The generated cDNA was diluted with sterile milliQ water. *Real Time PCR*. In each PCR tube 6 μl of cDNA was mixed with 0.375 μl of a 10 μM solution of each primer and 7.5 μl of 2× SYBR Green PCR Master Mix, to a final volume of 15 μl with sterile milliQ water (See Table 1 for primers sequences). Duplicates were prepared for each sample. Real-time PCR was performed on the thermal cycler Rotor-Gene Q (Qiagen, Venlo, Limburg, Netherlands) according to the following protocol: Initial denaturation 10 min at 95 °C, amplification (45 cycles) with denaturation 15 s at 95°C, annealing 30 s at 60°C and extension 30 s 70°C. To confirm the presence of a single product electrophoresis and a melting curve of the DNA was always made covering the range of 50-95°C. Relative gene expression levels were quantified using the comparative DCT method. This method normalized CT values of the detected gene to the average of that of the GAPDH endogenous control gene and calculated the relative expression values as fold changes of the control group (BMDM-WT without stimuli), which was set at 1. RT-PCRs were run in triplicate for each mouse.

### Nitric Oxide production assay

The concentration of nitrites in supernatants from BMDM cultures were obtained as an indirect measurement of NO accumulation using the Griess reaction^29^. 1×10^6^ BMDM were cultured in a 48-well plate for 22 h in the presence of LPS+IFN_γ_ or IL-4 at 37°C with 5% CO_2_. Subsequently, 100 μL of supernatant were placed in a 96-well flat-bottomed microplate and 100 μL of a mixture of 10 mg/ml sulfanilamide and 1mg/mL N-(1-naphthyl)-ethylenediamine dissolved in 2.5% H_3_PO_4_ and incubated for 10 min at room temperature in the dark. The optical density was then measured at a wavelength of 550 nm. The values obtained were expressed as nitrite (μM) concentration and extrapolated using a standard sodium nitrite curve.

### Superoxide production assay (O_2_^−^)

The production of O_2_^−^ by BMDM was evaluated by exposing the cells to Nitroblue Tetrazolium Chloride (NBT), solubilizing the blue formazan and quantifying its absorbance. 250.000 BMDM were cultured in a 96-well plate for 22 h in the presence of LPS+IFN_γ_ or IL-4 at 37°C with 5% CO_2_. Subsequently, supernatants were discarded by inversion and cells were incubated with a solution of 1mg/mL NBT diluted in PBS and filtered. After 30 min of incubation in an oven at 37°C, the cells were fixed with 100 L of 70% ethanol for 5 min. Finally, the blue precipitate was dissolved with 100 μL of KOH 2M and 100 μL of dimethyl sulfoxide and the optical density -OD- was determined at 655 nm. Higher optical density measured means higher O_2_^−^ production^30^.

### Whole-blood ROS levels measured by Electron Paramagnetic Resonance (EPR)

Trunk blood samples were harvested into heparinized tubes and processed for EPR analysis by using the spin probe, 1-hydroxy-3-carboxy-pyrrolidine (CPH). This EPR-silent cyclic hydroxylamine is oxidized to its derivative EPR-detectable nitroxide radical (CP radical), whose intensity is indicative of ROS formation. CPH is partially cell-permeable providing both extracellular and intracellular ROS formation. Moreover, it is not specific of a singular oxidant, but it is suitable to screen the totality of ROS produced in biological samples. Briefly, 100 μl of whole blood were incubated for 20 min at 37°C with CPH (1mM). Thereafter, samples were drawn up into a gas-permeable Teflon tube, folded two times, inserted into a quartz tube and fixed to the EPR cavity. Spectra were acquired 23 min after the addition of the spin probe at room temperature. Three independent experiments were performed and the values obtained were normalized for the proper statistical analysis.

### Statistical analysis

The results are expressed as mean ± SEM. Independent variables were analyzed using the t-test or two-way analysis of variance (ANOVA). Whenever ANOVA indicated significant effects (p<0.05), a Tukey *Post hoc* test was carried out. In all cases, the assumptions of the analysis of variance (homogeneity of variance and normal distribution) were attained. In all statistical analysis, a p<0.05 was considered to represent a significant difference between groups. All the analysis was performed using the software GraphPad Prism version 7.0 (La Jolla, California USA).

## RESULTS

### MeCP2 truncation in BMDM induces differential response to polarizing stimuli

In order to assess the ability of MeCP2^308/y^ (MUT) and wild-type WT bone marrow precursors to differentiate into macrophages, precursor cells were cultured for 6 days in the presence of M-CSF and the percentages of CD11b^+^CD45^+^ cells were analyzed by flow cytometry. We found that 96-98% of the cells in cultures from both genotypes were CD11b^+^CD45^+^, indicating a high level of purity (data not shown). Therefore, MeCP2 truncation did not affect the process of differentiation to macrophages under standard conditions.

Next, we sought to compare the immune response in WT and MUT-BMDM under polarizing conditions. For this purpose, we stimulated BMDM with either LPS+IFN_γ_ (classical activation, M1 profile) or IL-4 (alternative activation, M2 profile) and 4 h later, we assessed the expression levels of prototypical genes of each profile using real time RT-PCR.

We found that for both genotypes, the relative level of expression for all the M1 genes analyzed (TNFα, IL-1β, IL-6 and iNOS) was significantly increased in LPS+IFN_γ_-activated BMDM in comparison to basal levels (Fig. 1a-d). However, MUT-BMDM under this stimuli showed higher level of expression of TNFα compared to stimulated WT-BMDM (Fig. 1a). On the other hand, when WT-BMDM were stimulated with IL-4, we found significant increase in the expression of all M2 genes analyzed (FIZZ, CD206, ARG-1 and IL-10) in relation to the control group (Fig. 1e-h). Conversely, IL-4-stimulated MUT-BMDM showed significantly lower expression of FIZZ1 compared to stimulated WT-BMDM (Fig. 1e). Moreover, MUT-BMDM expressed lower levels of IL-10 and the surface marker CD206 than WT-BMDM, regardless of the stimulus (Fig. 1f, h). These results suggest that MeCP2 mutation alters the immune profile under polarizing conditions, resulting in an exacerbated pro-inflammatory setting and a deficient anti-inflammatory response.

**FIGURE 1.**
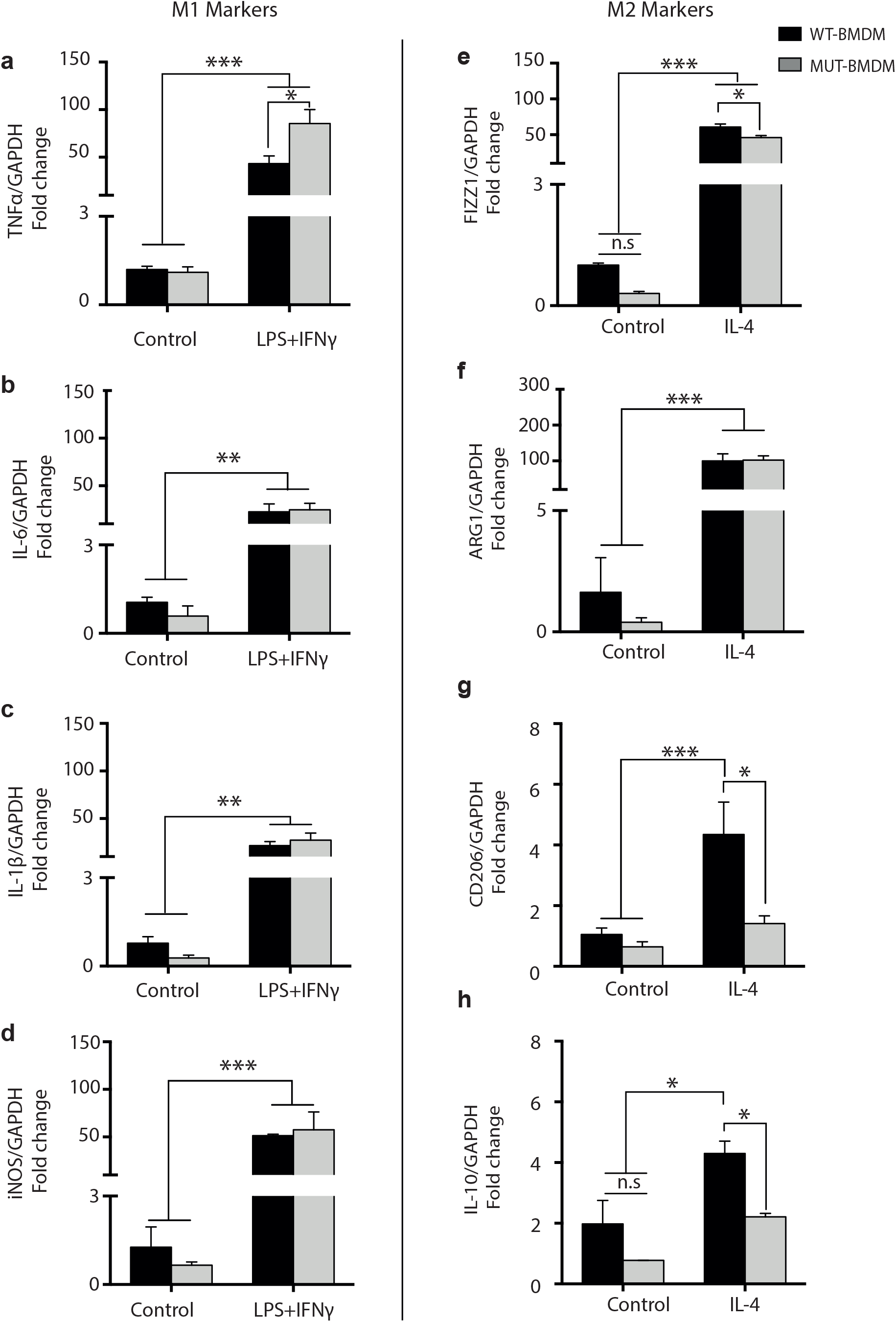
Relative mRNA abundance of prototypic genes in M1 and M2 BMDM. BMDM were stimulated *in vitro* for 4 h with LPS+IFN_γ_ or IL-4. For all genes analyzed (a-h) the fold change of expression was calculated using the 2^−ΔΔCT^ method. GAPDH was used as the housekeeping gene in all cases. **a-d)** For all M1 markers, the gene expression increased in response to LPS-IFN_γ_ in both WT and MUT-BMDM, and in this last group, a significant higher expression in TNFα was found compared to WT under stimulation (a). **e-h)** For M2 markers, the gene expression between control and stimulated BMDM in both, WT and MUT cultures, was significantly higher for FIZZ1 and ARG-1 genes (e,f). The analysis of gene expression of CD206 and IL-10 showed a significant increase in expression under IL-4 stimulation in WT-BMDM (g, h). Accordingly, MUT-BMDM showed significantly lower expression of both genes compared to WT-BMDM in response to IL-4. Data are presented as media ± SEM. Two-way ANOVA and *post hoc* Tukey test were performed. *p<0.05; **p<0.001 ***p<0.0001 (n per experimental group=4).

### MeCP2 truncation increases the level of reactive oxidizing species (ROS) in BMDM and in MeCP2^308/y^ mice

The production of reactive oxidizing species (ROS) actively participates in M1 responses acting as an effector molecule against pathogens^5^. However, if the production of these radicals is excessive, it can lead to protein oxidation, lipid peroxidation, through the increased production of superoxide radical 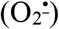, hydrogen peroxide (H_2_O_2_), hydroxyl radical (^•^OH) and peroxynitrite, all oxidants able to damage biomolecules^5^. In MeCP2-null, MeCP2^308/Y^ mice and RTT patients, it has been reported a persistent oxidative stress status^6,7^. To understand if the deficit of MeCP2 in macrophages could affect ROS generation, BMDM were stimulated with either LPS+IFN-_γ_ or IL-4 for 22 h and the 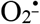 and the nitric oxide (NO) amounts were quantified in the culture supernatant. NO is a well-known powerful free radical with cytotoxic effects during inflammatory responses. We found that regardless the stimulus MUT-BMDM showed higher levels of 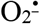 compared to WT-BMDM (Fig. 2a). Low levels of NO, measured as nitrite concentration, were detected in WT- and MUT-BMDM before and after stimulation with IL-4 (Fig. 2b). Conversely, when BMDM were stimulated with LPS+IFN_γ_, the nitrites accumulation was up to 12 times higher than the control and IL-4 stimulated groups (Fig. 2b). Since we observed that iNOS expression was similar in WT and MUT-BMDM after LPS+IFN_γ_ stimulation (Fig. 1d), our results suggest that the immune response mediated by iNOS and the consequent NO production upon M1-activation is not directly influenced by MeCP2.

**FIGURE 2.**
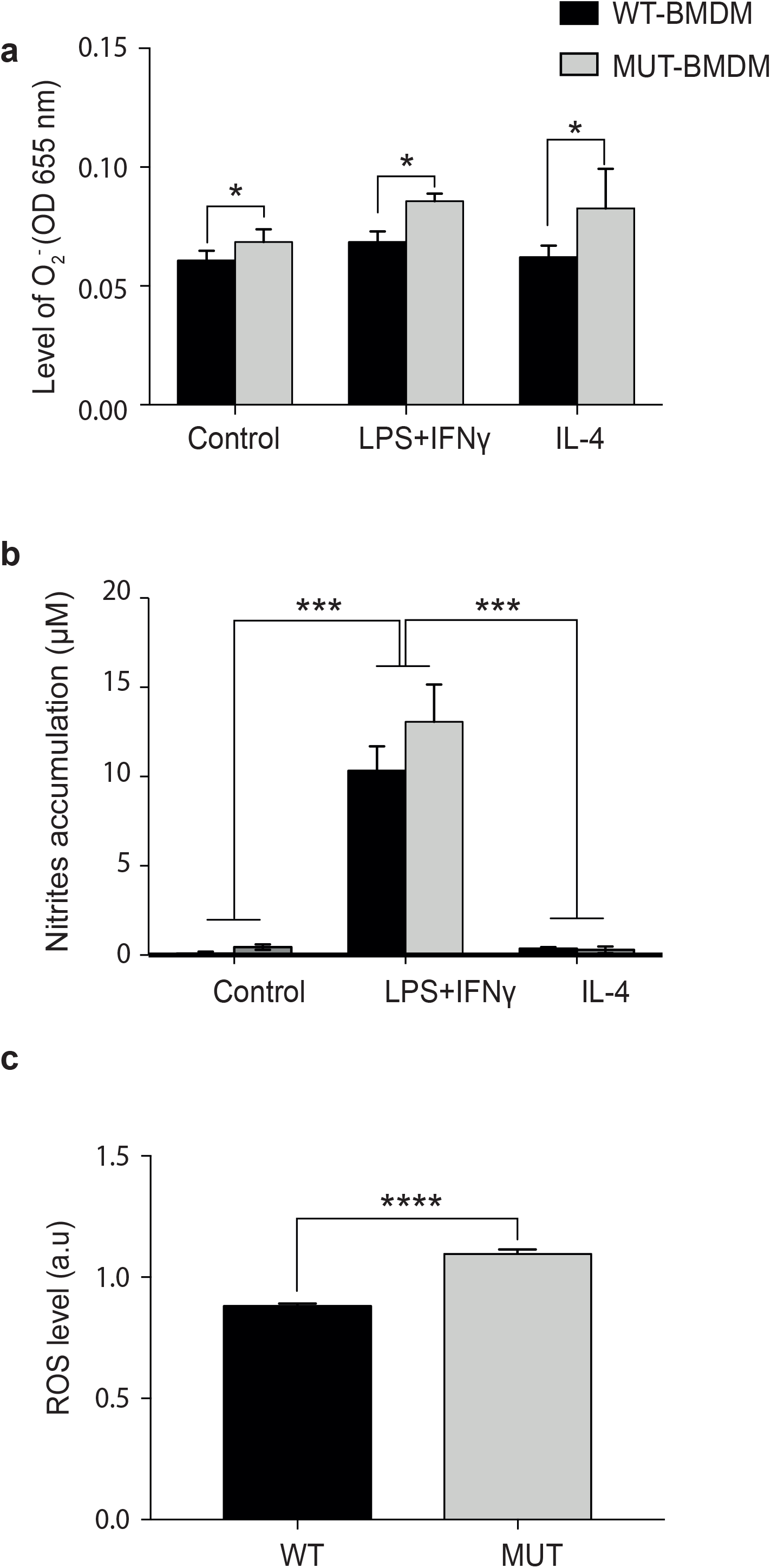
Level of production of M1 effector molecules in BMDM and mice. **a)** The graph shows the level of optic density (OD) measured at 655 nm as an indirect measurement of O2-production by BMDM. Each treatment, including controls, was carried out in triplicates. MUT-BMDM produced higher O_2^−^_ levels compared to WT-BMDM. Data are presented as media ± SEM. Two-way ANOVA was performed. *p<0.05. (n per experimental group= 4). **b)** The accumulation of nitrites (μM) was detected in culture supernatants as an indirect measurement of NO production by BMDM stimulated with LPS+IFN_γ_ or IL-4 for 22 h. Each treatment, including controls, was carried out in triplicates. No differences were found in the NO production between WT-BMDM and MUT-BMDM. Two-way ANOVA was performed. ****p<0.00001. (n per experimental group= 4). **c)** 12-weeks old mice were sacrificed and whole-blood was obtained. The level of ROS was determined through electron paramagnetic resonance (EPR). Values obtained in three independent experiments were normalized. MUT mice showed significantly higher levels of systemic ROS compared to WT animals. t-Student test was performed. **** p<0.0001. (n per experimental group=8-10).

Given this result we next aimed to establish the systemic level of ROS in these animals. For this, whole blood from WT and MUT mice was isolated and the level of ROS was obtained by electron paramagnetic resonance (EPR). We found that MUT mice produced significantly higher whole-blood ROS levels compared to WT (Fig. 2c). This result suggest that MeCP2 is an important factor in maintaining redox homeostasis under basal conditions, even in the absence of an inflammatory or oxidative stimulus.

## DISCUSSION

While many studies have addressed the role of MeCP2 in the nervous system, only a few have addressed the contribution of immune dysfunctions to the pathogenesis of RTT^3^. Macrophages are crucial effector cells able to orchestrate both innate and adaptive immune responses and they are indispensable for tissue homeostasis^8,9^. Since MeCP2 is a ubiquitous transcription factor and previous work have established that MeCP2 affects myeloid cells^3^, we sought to evaluate the potential role MeCP2 on macrophage polarization, in the context of an active immune response. For this purpose, we used cultures of BMDM isolated from MeCP2^308/y^ mice, a RTT mouse model that expresses a truncated form of MeCP2 and presents aberrant oxidative brain damage and peripheral inflammation^10^.

First, we were able to determine that the process of differentiation from bone marrow precursors to macrophages was not directly influenced by MeCP2 mutation according to the expression of specific markers CD45+CD11b+. Similar results were found in previous work using MeCP2-*null* Ly6c^high^ monocytes, which displayed similar kinetics of differentiation to macrophages and proliferative capacity as the WT^3^.

In response to LPS+IFN_γ_ (M1 stimuli), both WT and MUT-BMDM produced increased levels of prototypic M1 genes. However, MUT-BMDM expressed significant higher levels of TNFα transcript; this finding was similar to other reports using microglial and macrophages from MeCP2-*null* models and in RTT patients^3,11–13^. Interestingly, we found that none of the genes analyzed showed differences in expression between WT and MUT macrophages under baseline conditions indicating that immune activation is necessary for MeCP2 to modulate pro-inflammatory responses.

Although M1 macrophage activation converges on the NF-κB pathway to generates NO, and NF-κB pathway could be directly regulated by MeCP2 in peripheral blood mononuclear cells (PBMC) and splenocytes^13^, in the present study we found no differences in either NO production or iNOS expression levels between WT-BMDM and MUT-BMDM after LPS+IFN_γ_ stimulation. Our results support the idea that the role of MeCP2 on the expression of target genes is highly dependent on the cell type, the stimulation and the context (*in vivo* vs. *in vitro*)^15^.

ROS production is an important component of the M1 response, and are involved in several vital processes; however, excessive generation or insufficient detoxification generates a state of oxidative stress^16^. Here, we found higher basal levels of intracellular O_2^−^_ in MUT-BMDM, indicative of oxidative stress, as reported in patients and in MeCP2-*null* models^7,17–19^. In addition, we determine that ROS levels were significantly higher in whole blood of MUT mice. Therefore, we propose that MUT macrophage response may actively contribute to the imbalance observed in systemic ROS production and regulation. In RTT patients, it has been shown that the activity of enzymes that maintain redox balance, such as superoxide dismutase, is decreased and accompanied by increased lipid peroxidation^5^. Altogether, the implications of our results suggest that in the context of MeCP2 mutations, the exacerbated inflammatory profile could be maintained by a mechanism that feeds itself via the generation of oxidative imbalance; in this scenario macrophages could be affecting directly the tissue homeostasis *in vivo.*

Regarding anti-inflammatory and regulatory immune responses, other authors showed that the global response to glucocorticoids was increased in peritoneal macrophages and microglia from MeCP2-*null* mice, indicating that MeCP2 would act as a repressor of this pathway^3^. However, to our knowledge, the present work is the first study that has focused on characterizing M2 response in MeCP2-deficient macrophages. IL-4-stimulated MUT-BMDM showed lower expression levels of FIZZ1 and a failure to increase the expression of M2 prototypic genes IL-10 and CD206. In macrophages, miR-124 is essential for the induction and maintenance of the M2 phenotype^20^; interestingly, miR-124 is regulated by MeCP2 in CD4+ T cells^21^. These data indicate that the regulation given by MeCP2 to maintain the M2 profile in macrophages could be complex; future experiments will address the possible actors involved in this regulation. Concerning IL-10, evidence from other cell types (astrocytes and muscle cells) indicate that MeCP2 participates in the regulation of IL-10 responses by indirect mechanisms (i.e. via miR-181)^15,22^. We propose that it could also be the case for MeCP2-mediated IL-10 regulation in macrophages.

FIZZ1 is induced under allergic lung inflammation and it stimulates the expression of α-SMA (smooth muscle actin) and collagen in lung fibroblasts^23^. In turn, α-SMA is indirectly regulated by MeCP2 and in fact, MeCP2-*null* mice show less lung fibrosis^24^. Since this factor has been recently described, little is known about the role of FIZZ1 in the context of MeCP2 mutations. Understanding the interaction MeCP2/FIZZ-1 may have important therapeutic implications for patients with RTT, due to the recurrent pulmonary conditions they present^25^.

In summary, we can confirm that MeCP2 is an important regulator of macrophage function by modulation of pro- and anti-inflammatory gene expression *in vitro* (Fig. 3). Future work will assess functional characterization of MeCP2-deficient macrophages, in order to evaluate the potential involvement in Rett Syndrome pathogenesis. Given some discrepancies in the expression of genes analyzed in this work and previous studies^13,14,26,27^, we emphasize that the influence of MeCP2 would be dependent on different factors, such as the total absence of MeCP2 vs. its presence as a mutated protein, cell type, stimulus and context (*in vitro* vs *in vivo*). Further research should be oriented to understand the consequences MeCP2 mutations on immune imbalance in the whole organism, an essential information to consider possible therapeutic interventions in MeCP2-related disorders.

**FIGURE 3.**
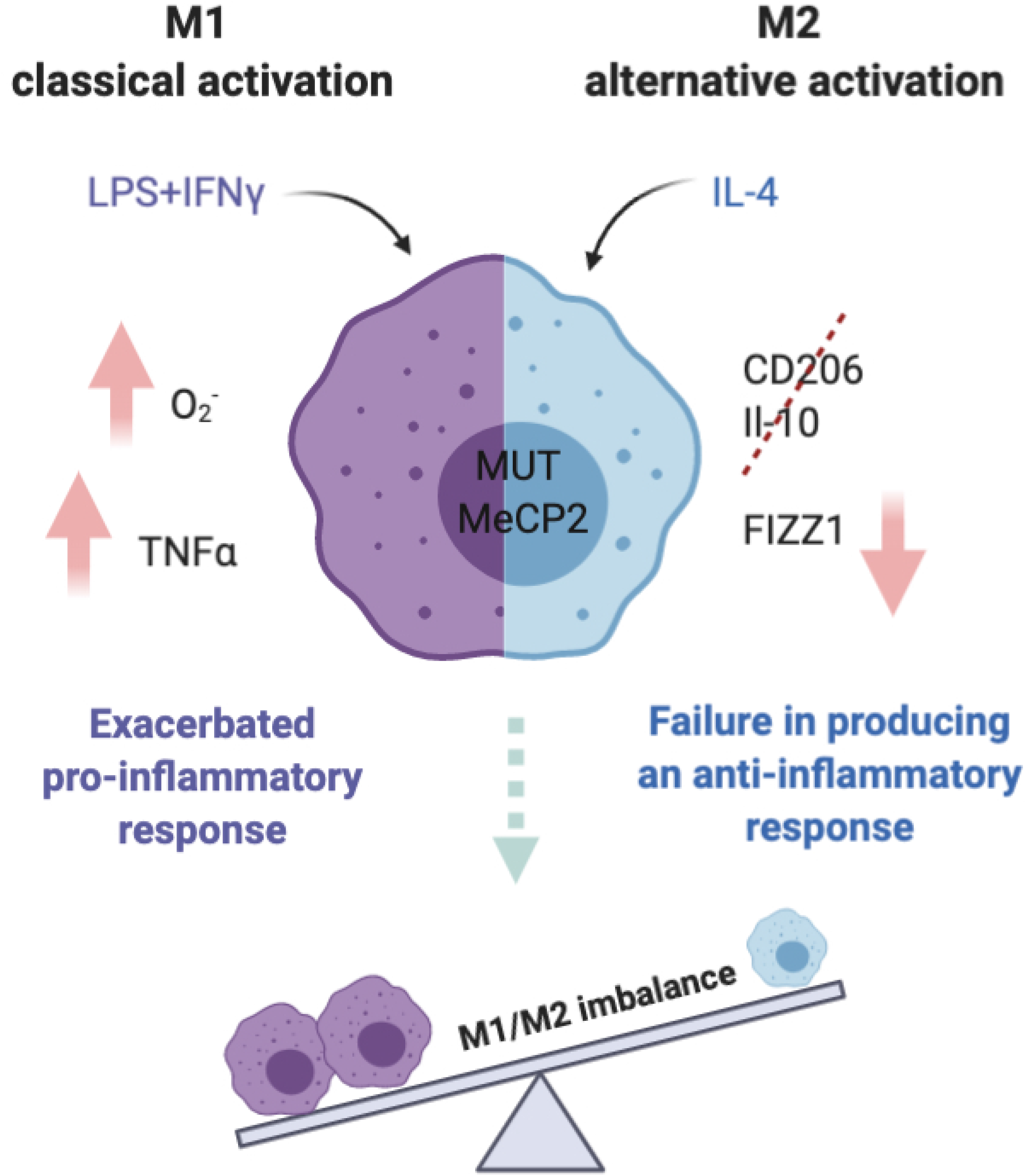
Schematic summary of Mecp2^308/y^ BMDM response to polarizing stimuli. MeCP2 truncation leads to increased ROS production under basal conditions. Under M1 stimulation, MeCP2 truncation induces an exacerbated pro-inflammatory response by expressing increased levels of TNFα transcripts. Under M2 activation, MUT-BMDM show deficient anti-inflammatory response given by poor CD206, IL-10 and FIZZ1 upregulation. Our results indicate that MeCP2 plays a role in the establishment of macrophage polarization in the context of immune activation.

## Acknowledgments

This work was supported by grants FONCyT-PICT-2013-106 and Secyt-Universidad Nacional de Córdoba, Argentina to A.L.D. The study was *facilitated* by a grant of the Italian Ministry of Health (#GR-2018-12366210) to B.D.F. MIZ was supported with a PhD fellowship financed by CONICET and received a travel grant from ISN. ALD is a member of the Scientific Career of CONICET.

## Author Contributions

MI Zalosnik performed experiments, collected the data, analyzed and interpreted the data, wrote the first draft. B De Filippis, D Pietraforte, R De Simone and G Laviola provided key reagents, analyzed and interpreted the data. AL Degano conceived and designed the study, supervised the study, wrote the paper, and approved the final version of the manuscript. All authors contributed to revise the manuscript critically and approved the submitted version.

## Notes

### Competing Interest Statement

The authors have declared no competing interest.

