## Supplementary material for "Mecp2 Deficiency Alters M1/M2 Gene Expresion in Bone Marrow-Derived Macrophages Upon Stimulation": Table 1

**Table 1.** Summary of the primer sequence used for Real Time RT-PCR and product sizes obtained.

| Target gene | Primers Sequences | | Product size (bp) |
|---|---|---|---|
| | pForward | pReverse | |
| **IL-1β** | CAGGCAGGCAGTATCACTCA | TGTCCTCATCCTGGAAGGTC | 86 |
| **TNFα** | AGCCCCCAGTCTGTATCCTT | GGTCACTGTCCCAGCATCTT | 113 |
| **IL-6** | CCGGAGAGGAGACTTCACAG | TCCACGATTTCCCAGAGAAC | 102 |
| **iNOS** | GTTCTCAGCCCAACAATACAAGA | GTGGACGGGTCGATGTCAC | 127 |
| **IL-10** | CATGGGTCTTGGGAAGAGAA | AACTGGCCACAGTTTTCAGG | 117 |
| **FIZZ1** | TGGCTTGCGAGACGTAGAC | ACCCAGTAGCAGTCATCCCA | 108 |
| **ARG-1** | TGGCTTGCGAGACGTAGAC | GCTCAGGTGAATCGGCCTTTT | 160 |
| **CD206** | CAAAAACTGACTGGGCTTCC | GCCCTTGATTCCAAAGAGTG | 101 |

IL-1β interleukin 1-beta; TNFα, tumor necrosis factor alfa; IL-6, interleukin 6; iNOS, inducible nitric oxide synthase; IL-10, interleukin 10; FIZZ1, found in inflammatory zone 1; ARG-1, arginase 1; CD206, cluster of differentiation 206.
